# miR-26a regulates extracellular vesicle secretion from prostate cancer cells via targeting SHC4, PFDN4 and CHORDC1

**DOI:** 10.1101/646380

**Authors:** Fumihiko Urabe, Nobuyoshi Kosaka, Yurika Sawa, Tomofumi Yamamoto, Yusuke Yamamoto, Kagenori Ito, Takahiro Kimura, Shin Egawa, Takahiro Ochiya

**Author notes:** **Corresponding author:** Nobuyoshi Kosaka, PhD, Department of Molecular and Cellular Medicine, Tokyo Medical University, Tokyo, Japan.

## Abstract

Extracellular vesicles (EVs) are known to be involved in intercellular communication during cancer progression; thus, elucidating the detailed mechanism will contribute to the development of a novel strategy for EV-targeted cancer treatment. However, the biogenesis of EVs in cancer cells is not completely understood. MicroRNAs (miRNAs) regulate a variety of physiological and pathological phenomena; thus, miRNAs could regulate EV secretion. Here, we performed high-throughput miRNA-based screening to identify the regulators of EV secretion using an ExoScreen assay. By using this miRNA-based screening, we identified miR-26a, which was reported as a tumor suppressive miRNA, as a miRNA involved in EV secretion from prostate cancer (PCa) cells. In addition, we found that the SHC4, PFDN4, and CHORDC1 genes regulate EV secretion in PCa cells. Suppression of these genes by siRNAs significantly inhibited the secretion of EVs in PCa cells. Furthermore, the progression of PCa cells was inhibited in an in vivo study. On the other hand, injection of EVs isolated from PCa cells partially rescued this suppressive effect on tumor growth. Taken together, our findings suggest that miR-26a regulates EV secretion via targeting SHC4, PFDN4, and CHORDC1 in PCa cells, resulting in the suppression of PCa progression.

## Introduction

Extracellular vesicles (EVs) include a wide variety of small membrane-bound vesicles that are actively released from almost all types of cells^1^, and play important roles in intercellular communication. EVs transfer of functional molecules, including miRNAs, mRNAs, proteins and lipids into the recipient cells. Through the transfer of these contents, EVs have been demonstrated not only function in normal physiological processes ^2^, but also be associated with the pathogenesis of various diseases. Especially in cancer field, number of studies have shown that EVs play important roles in tumor progression ^3^. Indeed, in prostate cancer (PCa), some reports have shown that EVs contribute to drug resistance or progression of metastasis^4 5 6^.

Recently several reports have shown the potential that a reduction in cancer-derived EVs shows therapeutic value by inhibiting cancer proliferation and dissemination ^3^. For instance, HER2 expressed on the surface of breast cancer-derived EVs has been shown to interfere with therapy and is associated with cancer progression ^7^. In addition, Marleau et al. described a therapeutic strategy for the removal of circulating EVs by developing a hemofiltration system to capture HER2-positive EVs ^8^. Furthermore, we recently showed that the administration of antibodies against human-specific CD9 and CD63, which are enriched on the surface of EVs, significantly decreased metastasis in a human breast cancer xenograft mouse model ^9^. These reports provide promising evidence that the inhibition of circulating EVs could be a novel strategy for cancer treatment. EV secretion from cancer cells was higher than that from normal cells, suggesting that cancer cells have a gene regulation network to promote EV production and/or EV secretion ^10^. Thus, understanding this regulatory network will have significant therapeutic value in cancer. However, despite significant advances in understanding the role of EVs in cancer progression, investigation of the biogenesis of EVs in cancer cells remains obscure. Therefore, the identification of the mechanisms of EV biogenesis will have significant therapeutic potential in cancer.

MicroRNAs (miRNAs) are small noncoding RNAs of 20-25 nucleotides in length that post-transcriptionally regulate the expression of thousands of genes, and a growing body of evidence has shown that miRNAs are the key regulators of several biological processes. Importantly, miRNAs are closely associated with tumorigenesis and several stages of metastasis ^11^. In noncancer cells, miRNAs systematically regulate RNA molecular networks; however, in cancer cells, aberrantly expressed miRNAs disrupt the otherwise tightly regulated relationship between miRNAs and mRNAs, leading to progression and metastasis. As shown previously, EV is involved in cancer progression; thus, we hypothesized that miRNAs could regulate EV secretion in cancer cells.

In this study, we found a novel mechanism of EV secretion in PCa cells by investigating miRNAs that are involved in EV secretion. To perform screening of nearly 2000 species of miRNAs, we used our established EV detection method, ExoScreen, which can directly detect EVs in conditioned medium based on an amplified luminescent proximity homogeneous assay ^12^. We comprehensively screened miRNAs using a miRNA library and found that miR-26a, which was reported as a tumor suppressive miRNA in PCa ^12 13^, negatively regulates EV secretion in PCa cells. In addition, we identified three target genes that were involved in EV secretion in PCa cells. Furthermore, reduced expression of miR-26a and upregulation of target genes were shown in PCa tumors compared with normal tissues. An in vivo study demonstrated that reduced expression of these three genes inhibited PCa tumor growth, and this change was partially rescued by the injection of EVs from PCa cells. These results suggest novel insight into miRNA-mediated tumor suppression through inhibiting EV biogenesis, which may provide novel approaches for PCa treatment.

## Results

### Establishment of a high-throughput compatible extracellular vesicle biogenesis assay

The PC3M cells were seeded in 96-well plates and transfected with each miRNA mimic. Twenty-four hours after transfection, the medium was changed to serum-free medium and then incubated for another 48 hours. After that, we collected the conditioned medium ^14^ to evaluate the EV level by ExoScreen, which can directly detect EVs based on an amplified luminescent proximity homogeneous assay using photosensitizer beads and two specific antigens residing on EVs ^12^ (Figure 1A). We confirmed that EVs derived from PC3M were CD9- and CD63-positive by immunoblotting (Figure 1B); therefore, we used CD9 to detect the change in EV secretion in this first screen. In addition, to exclude the effect of miRNAs on cellular proliferation, we performed a colorimetric MTS assay, as shown in Figure 1A. To assess the quality of transfection in each plate, several controls were used, and the effectiveness of siRNA controls on EV secretion was almost the same between the plates and validated the quality of transfection (Supplementary Figure 1A). The EV secretion was calculated by the ExoScreen assay and MTS assay, and the values were evaluated as the fold change relative to the negative control.

**Figure 1.**
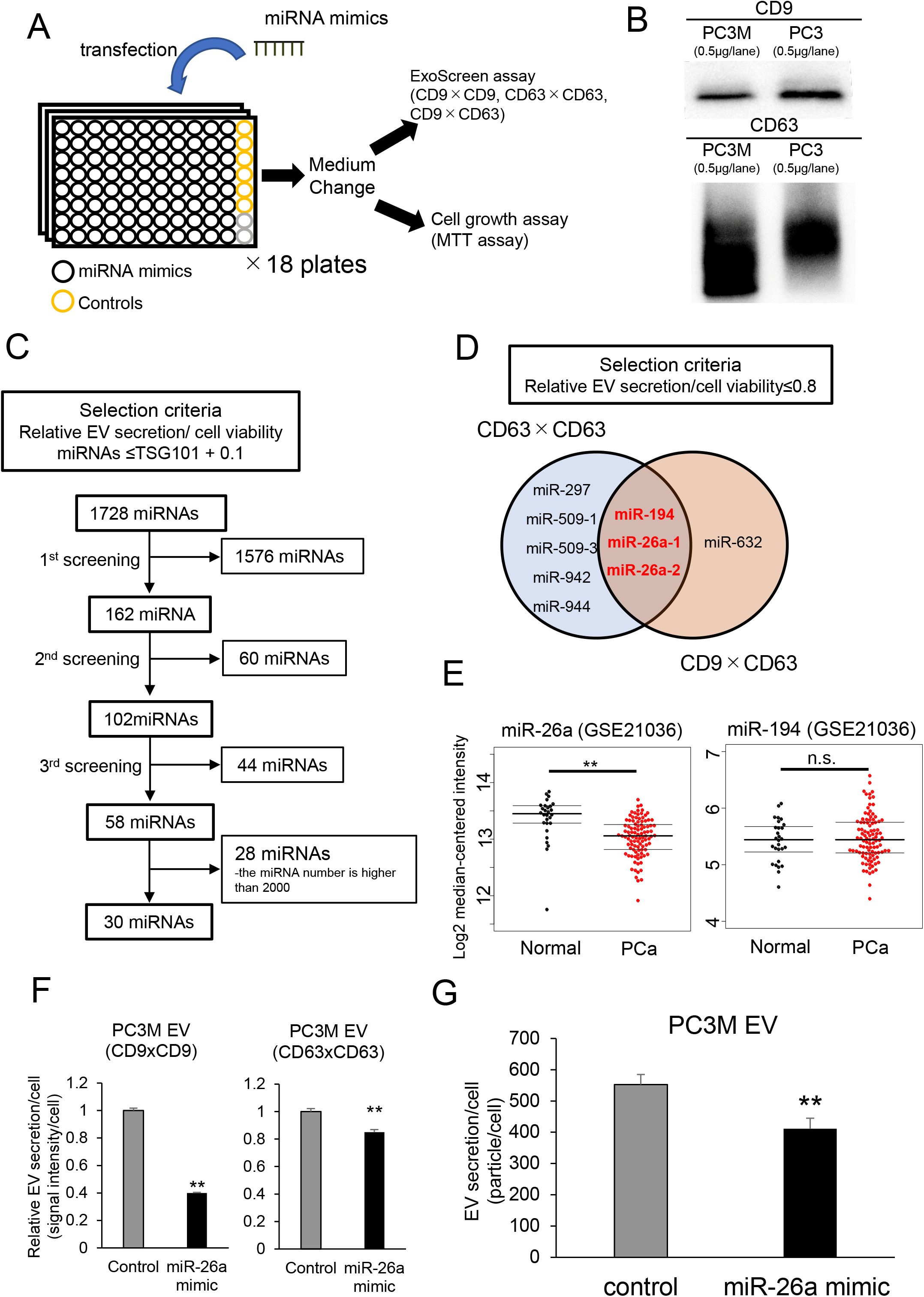
Screening of miRNAs regulating EV secretion in prostate cancer. A. Schematic illustration of a high-throughput compatible extracellular vesicle (EV) biogenesis assay to detect EV biogenesis-regulating miRNAs. B. Immunoblot analysis of the conventional EV markers CD9 and CD63 on EVs from PCa. C. Flow diagram of miRNAs used for selecting candidate miRNAs. D. Venn diagram showing miRNAs that suppress EV secretion. The miRNAs whose relative EV secretion/cell viability was lower than 0.8 were selected in each assay. The secretion of EVs was evaluated by ExoScreen, and the cell viability was measured by the MTS assay. E. Expression levels of miR-26a and miR-194 in prostate cancer clinical specimens (GSE21036). **, p<0.01; and n.s., not significant. F. The effect of the miR-26a mimic on EV secretion per PC3M cell. The amount of secretion of EVs per cell was evaluated by the signal intensity of ExoScreen per cell. The values are depicted as the fold change relative to the non-specific miRNA mimic (control). The values are the mean±SE (n=3). **, p<0.01. G. The effect of the miR-26a mimic on EV secretion per PC3M cell. The amount of EV secreted per cell was evaluated using a nanoparticle tracking system. The values are the mean±SE (n=3). **, p<0.01.

### Quantitative high-throughput analysis of candidate miRNAs in prostate cancer cells

A miRNA mimic library was screened to investigate the modulatory effects of various kinds of miRNAs on EV biogenesis. We evaluated the effectiveness of each miRNA on the secretion of EVs by ExoScreen and cell proliferation by colorimetric MTS assays. We selected miRNAs according to the criteria shown in Figure 1C. We performed screenings three times and chose 58 miRNAs. After excluding miRNAs whose number was higher than 2000, we selected 30 miRNAs (Figure 1C). Next, to further validate our initial screening, we assessed the secretion of CD63-positive EVs and CD9 and CD63 double-positive EVs by ExoScreen in these 30 miRNAs (Figure 1A). In this set, we selected miRNAs that showed the relative value of EV secretion/cell viability, evaluated by the ExoScreen assay and MTS assay, which was lower than 0.8. Since the relative value of EV secretion/cell viability by silencing TSG101, which is known to regulate the biogenesis of EVs ^15^, was 0.77, as evaluated by CD9 and CD63 double positive EVs (Supplementary Figure 1B), we expected that the miRNAs could suppress the secretion of EVs, similar to TSG101. Then, we chose miR-26a and miR-194 as candidate miRNAs to regulate EV secretion (Figure 1D). To select miRNAs that can clinically regulate EV secretion, we investigated a public database (GSE 21036). Principal component analysis (PCA) maps with 373 miRNAs suggested that the miRNA profiles differed between the PCa and normal adjacent benign prostate tissues (Supplementary Figure 2A). Additionally, as shown in a heat map displaying the 59 differentially expressed miRNAs, which were repressed more than 1.25-fold in prostate cancer tissue relative to normal adjacent benign prostate tissue and had a p-value less than 0.001, there were obvious differences in miRNA expression, including miR-26a, although we could not find a difference in the expression of miR-194 in prostate cancer tissue relative to normal adjacent benign prostate tissue (Figure 1E and Supplementary Figure 2B). These results suggest that miR-26a is involved in EV secretion of PCa. Furthermore, we confirmed that the particle number of EVs secreted by each PCa cell transfected with the miR-26a mimic was also decreased using ExoScreen and nanoparticle trafficking analysis (NTA) (Figure 1F, G and Supplementary Figure 2C, D). Therefore, we selected miR-26a for further or detailed analysis and investigated whether miR-26a regulates EV secretion in prostate cancer.

### Selection of candidate genes regulating extracellular vesicle secretion in prostate cancer cells

miRNAs are known to regulate hundreds of mRNA targets, providing global changes in the cellular phenotype of cells ^16^. To further elucidate the molecular mechanisms of miR-26a in EV secretion, we identified the target genes of miR-26a. We performed mRNA microarray analysis in PC3M and PC3 after the transfection of miR-26a mimic or control. For the genes that could be targeted by miR-26a picked up by TargetScan, we found that overexpression of miR-26a in prostate cancer cells downregulated 88 genes compared with the control cells by miRNA expression (Figure 2A). Then, to select genes regulating EV secretion, we performed high-throughput screening using ExoScreen again. The PC3M cells were seeded in a 96-well plate and transfected with each candidate siRNA of the 88 genes. Twenty-four hours after transfection, the medium was changed to serum-free medium for 48 hours of incubation. From the transfected PC3M cells, we collected the conditioned medium to evaluate the EV levels by ExoScreen and MTS assays (Figure 2B). We evaluated CD9-positive EVs and CD63-positive EVs by ExoScreen. The criteria of the selected genes are described in Figure 2C. The results of each screening are shown in Supplementary Figure 3. Finally, we identified four genes, SHC4, PFDN4, CHORDC1 and PRKCD, as candidate genes regulating EV secretion (Figure 2C).

**Figure 2.**
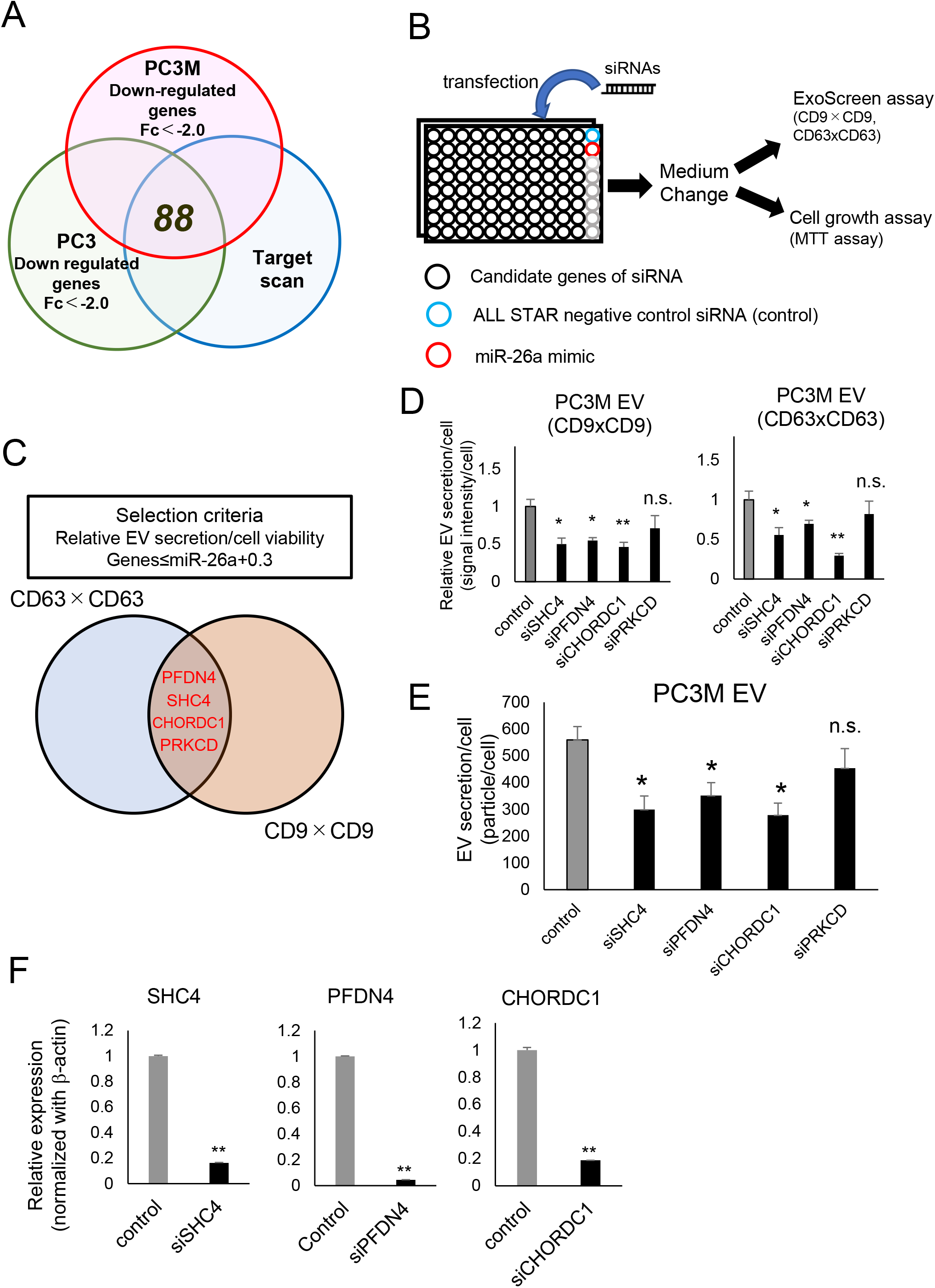
SHC4, PFDN4 and CHORDC1 are involved in miR-26a-mediated EV secretion. A. Venn diagram of predicted miR-26a targets (TargetScan) and transcripts that were experimentally repressed >2-fold by miR-26a overexpression in prostate cancer cells (PC3M or PC3) relative to control conditions. B. Schematic of the high-throughput compatible EV biogenesis assay to choose EV biogenesis-regulating genes. C. Venn diagram showing genes that suppress EV secretion evaluated by ExoScreen. The genes whose relative EV secretion/cell viability was lower than that of miR-26a plus 0.3 were selected in each assay. The secretion of EV was evaluated by ExoScreen, and the cell viability was measured by the MTS assay. D. The effect of siRNAs against candidate genes on EV secretion in PC3M cells. The EV secretion per cell was evaluated by the signal intensity of ExoScreen per cell. The values are depicted as the fold change relative to the negative control siRNA (control). The values are the mean±SE (n=3). *, p<0.05; **, p<0.01; and n.s., not significant. E. The effect of siRNAs against candidate genes on EV secretion per PC3M cell. The particle number of EVs was measured using a nanoparticle tracking system. The values are the mean±SE (n=3). *, p<0.05; and n.s., not significant. F. The effect of SHC4, PFDN4 and CHORDC1 siRNA on the mRNA expression level of each gene. β-actin was used as an internal control. Error bars represent the s.e. deduced by Student’s t-test (*P<0.05, **<0.01). n.s., no significant difference. The data are representative of at least three independent experiments. The values are the mean±SE (n=3). **, p<0.01.

### SHC4, PFDN4 and CHORDC1 regulate extracellular vesicle secretion in prostate cancer

Next, we confirmed the effect of these genes on the secretion of EVs derived from prostate cancer cells after treatment with siRNA for these genes. The EV levels secreted by each prostate cancer cell were decreased after transfection with the siRNAs of the three genes, SHC4, PFDN4 and CHORDC1, indicating that these genes are regulators of EV secretion (Figure 2D, E and Supplementary Figure 4A, B). The downregulation of the three genes in PCa cells transfected with siRNA of each gene was confirmed by qRT-PCR (Figure 2F). These results suggest that SHC4, PFDN4 and CHORDC1 could contribute to the upregulation of EV secretion in PCa.

### miR-26a suppressed extracellular vesicle secretion in prostate cancer cells by targeting SHC4, PFDN4 and CHORDC1

First, we confirmed that miR-26a suppressed the expression levels of SHC4, PFDN4 and CHORDC1 in PCa cells by qRT-PCR (Supplementary Figure 5A, B) and immunoblot analysis (Figure 3A). Then, to address whether miR-26a directly regulated these genes, we performed a luciferase reporter assay (Figure 3B). Ectopic expression of miR-26a significantly suppressed the luciferase activity of the wild-type SHC4, PFDN4 and CHORDC1 3’-UTRs but not their mutant 3’-UTRs (Figure 3C). These results provide experimental evidence that miR-26a can directly repress translation initiation of SHC4, PFDN4 and CHORDC1, and the downregulation of miR-26a promoted EV secretion. Additionally, using a public database (GSE6099), we investigated the expression levels of these genes in prostate cancer tissue. We confirmed that the expression levels of PFDN4 and CHORDC1 were significantly upregulated in prostate cancer tissue compared to normal tissue (Figure 3D).

**Figure 3.**
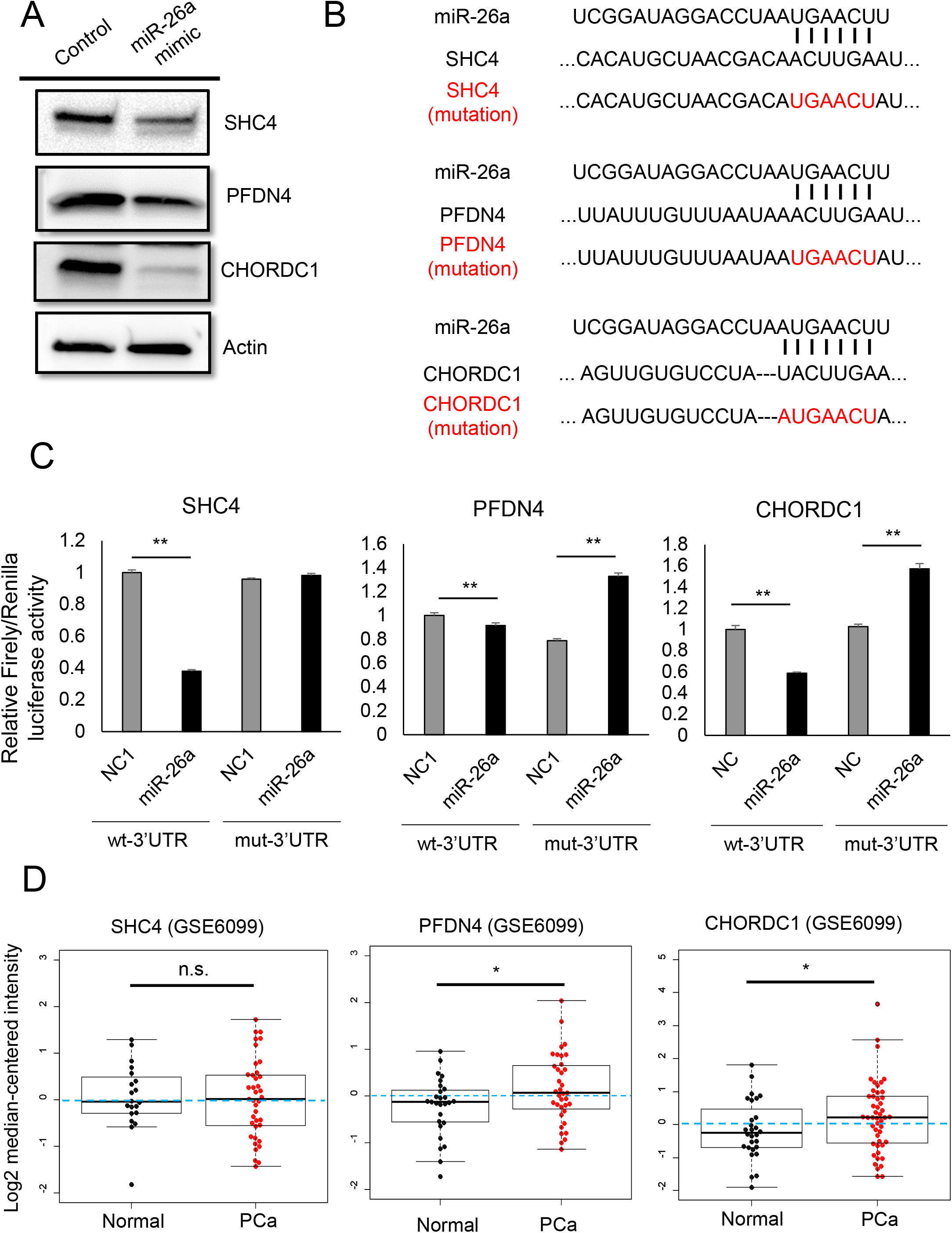
miR-26a directly regulates the expression levels of SHC4, PFDN4 and CHORDC1. A. Immunoblot analysis of PC3M cells transfected with nonspecific miRNA mimic (negative control mimic) or miR-26a mimic. B. Summary of miR-26a target sites and mutated sites (shown in red) in the 3’-UTRs of SHC4, PFDN4 and CHORDC1. C. Target validation of SHC4, PFDN4 and CHORDC1 was confirmed in the luciferase reporter assay. The values are depicted as the fold change relative to the negative control siRNA (control). The values are the mean±SE (n=3). **, p<0.01. D. Expression levels of SHC4, PFDN4 and CHORDC1 in prostate cancer and normal prostate tissue clinical specimens (GSE6099). *, p<0.05; and n.s., not significant.

### PFDN4, SHC4 and CHORDC1 regulate EV secretion and promote tumorigenesis in vivo

miR-26a was shown to suppress the tumor formation of prostate cancer; thus, this antitumor activity could be because of the suppression of EV secretion from prostate cancer cells by miR-26a.^13 17 18 19^. To confirm the role of these genes targeted by miR-26a in prostate cancer-derived EVs, we performed in vivo experiments. Initially, we established a PC3M cell line with stable SHC4, PFDN4 or CHORDC1 depletion by using each short-hairpin RNA and evaluated the common aspects of tumorigenesis. Depletion of these genes repressed EV secretion (Figure 4A). We assessed the effect of SHC4, PFDN4 and CHORDC1 downregulation in this model and found that mice bearing PC3M xenografts with depletion of these genes had smaller tumors that weighed less than those of control mice (Figure 4B, C). In addition the tumor tissues of nude mice injected with PC3M-derived EVs showed partially rescued tumor size and weight (Figure 4D and Supplementary Figure 6). The above data suggest a signaling network that links miR-26a with its targets SHC4, PFDN4, and CHORDC1 and demonstrated the novel mechanism of miR-26a-regulated EV secretion in prostate cancer (Figure 5).

**Figure 4.**
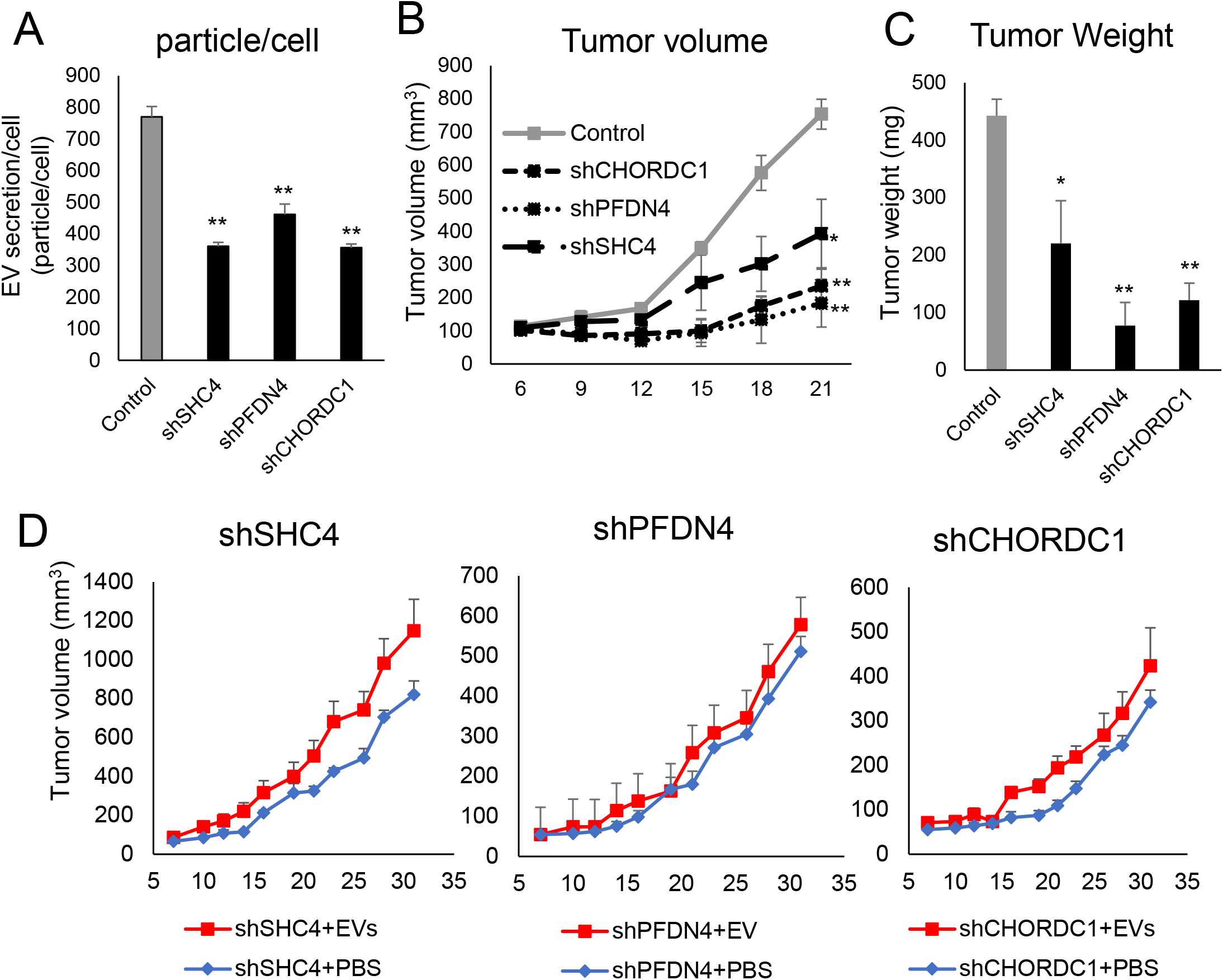
Downregulation of EV secretion inhibits cancer progression in vivo. A. Establishing the PC3M cell line with stable SHC4, PFDN4 and CHORDC1 depletion using short-hairpin RNAs and evaluation of EV secretion. The values are the mean±SE (n=3). **, p<0.01. B. The tumor volumes were measured every 3 days after tumor inoculation. The values are the mean±SE (n=3). *, p<0.05; and **, p<0.01. C. The tumor weights in nude mice at day 21 were determined. The values are the mean±SE (n=3). *, p<0.05; and **, p<0.01. D. The tumor volumes were measured every other day before the injection of EVs. The values are the mean±SE (n=6).

**Figure 5.**
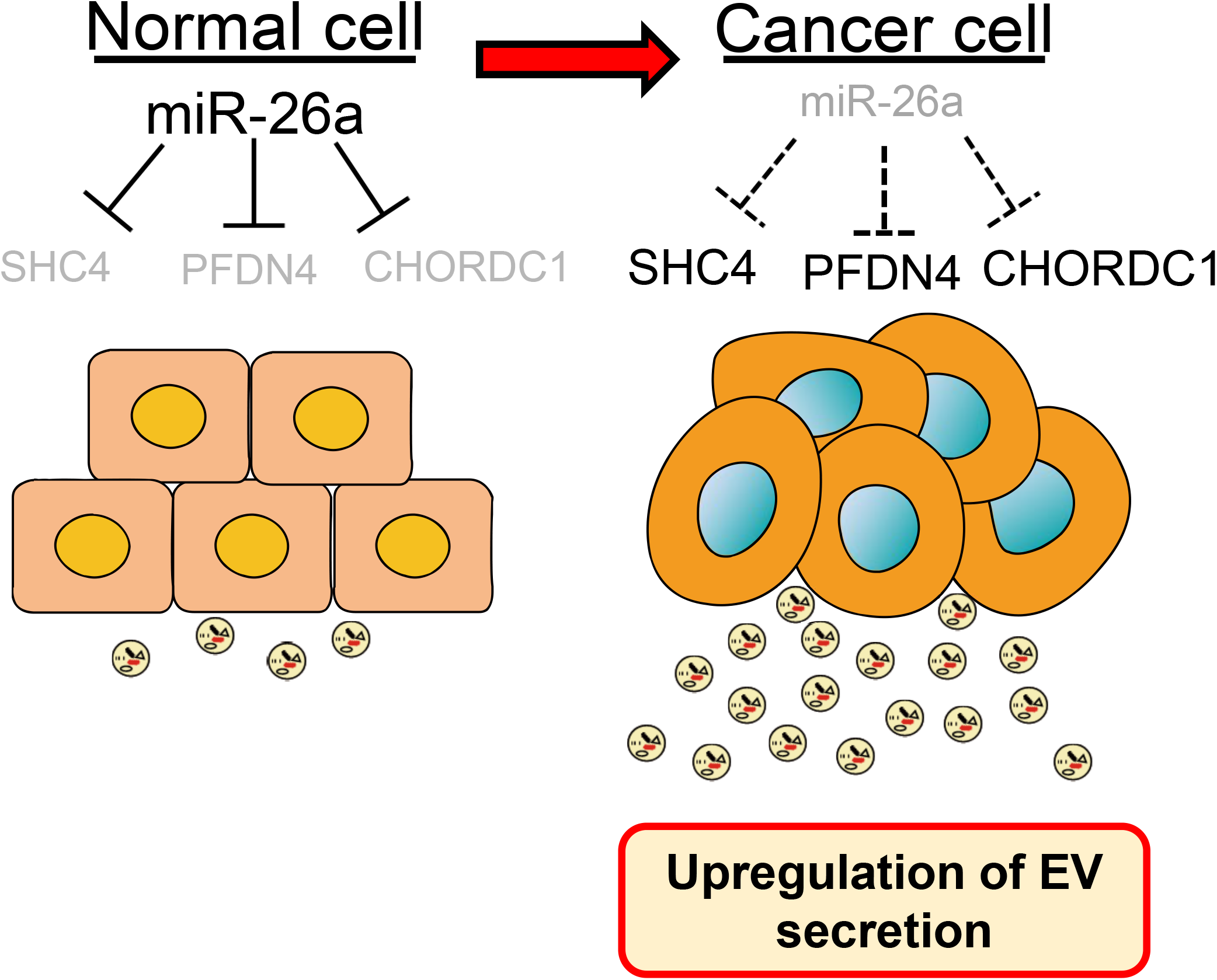
Schematic model of the regulation of EV secretion in prostate cancer.

## Discussion

Although EVs have been reported to modulate cancer progression for approximately ten years ^20^, the biogenesis of EV in cancer cells remains unclear. The endosomal sorting complex required for transport (ESCRT) machinery and their associated proteins, including TSG101 ^15^, Alix ^21^, and VPS4 ^22^, have been implicated in EV secretion. In addition, we showed that neutral sphingomyelinase-2 (nSMase2), which is required for the synthesis of ceramide, regulates EV secretion in breast cancer^10^. However, the downregulation of nSMase2 did not inhibit EV secretion in PCa ^23^. These data suggest that the biogenesis of EV secretion is different between each kind of cancer. In addition, to establish a novel therapeutic strategy for PCa by inhibiting EV secretion, the mechanism of PCa-specific EV secretion should be revealed.

In the present study, based on a comprehensive miRNA analysis, we showed that miR-26a and miR-194 regulate the secretion of EVs in PCa. We focused on miR-26a because the expression level of miR-26a was reduced in PCa tissue compared to normal prostate tissue. Previous studies have demonstrated the suppressive role of miR-26a in prostate cancer growth ^17 13 18 19^, and our study has reported the novel role of miR-26a. That is, miR-26a not only inhibits tumorigenesis but also prevents the secretion of EVs. As mentioned above, suppression of EV secretion could be a novel therapeutic strategy. Therefore, these data suggest that miR-26a represents a potential therapeutic target to treat the progression of prostate cancer by inhibiting EV communication.

miRNAs are known to regulate hundreds of mRNA targets, providing global changes in the cellular phenotype of cells ^16^. To investigate the target genes of miR-26a that suppress EV secretion, we performed high-throughput screening and selected four candidate genes, PRKCD, SHC4, PFDN4 and CHORDC1. Although the knockdown of PRKCD in PCa decreased the secretion of total EVs, it significantly decreased PCa cell proliferation. Therefore, the secretion of EVs by each cell was not decreased. PRKCD is implicated in the growth, migration and invasion of cancer cells, including PCa ^24^; therefore, downregulation of PRKCD in PCa cells was critical for cell viability.

We revealed that miR-26a directly targets SHC4, PFDN4 and CHORDC1, and downregulation of these genes suppressed the secretion of EVs in PCa. Although we did not report the precise mechanism of these genes in regulating EV secretion in the present study, several articles have suggested a relationship between EV biogenesis and these genes. SHC4 is one of the proteins of the Src homology and collagen family, ShcA, ShcB, ShcC and SHC4 (or ShcD), in chronological order of their discovery. Shc proteins are known to engage in the EGFR internalization process ^25^, and ShcD was reported to alter steady-state EGFR trafficking dynamics, which reduces cellular ligand sensitivity by recruiting the EGFR into juxtanuclear vesicles identified as Rab-11-positive endocytic recycling compartments ^26^. Rab-11 contributes to the endocytic pathway through transportation of the endocytosed cargo to the endocytic recycling compartment ^27^. In addition, Rab11 was found to be essential for EV secretion ^28^. These data suggest that ShcD may also affect the endocytic pathway and contribute to EV secretion. Morgana, which is coded by the CHORDC1 gene, binds to the Hsp90 chaperone protein, behaving as a co-chaperone ^29^. Recently, Lauwers et al. reported that Hsp90 directly binds and deforms membranes, thereby promoting the fusion of multivesicular bodies with the plasma membrane and the release of EVs ^30^. Therefore, CHORDC1 may support the secretion of EVs via stabilizing Hsp90. Compared with SHC4 and CHORDC1, much less is known about the function of PFDN4. The PFDN4 gene is a transcriptional factor regulating the cell cycle ^31^. In breast cancer, PFDN4 was found to play a role in cancer behavior; however, biological analysis remains to be performed ^32^. The precise mechanism by which these genes in EV secretion should be investigated in a future study; however, our data suggest, for the first time, that SHC4, PFDN4 and CHORDC1 contribute to EV secretion in prostate cancer and could be novel therapeutic targets.

Additionally, in vivo experiments showed that knocking down these genes decreased tumor progression at the primary site. Although we could not confirm that the tumor suppressive effect was only caused by the inhibition of EV secretion, the tumor xenografts of nude mice injected with PC3M-derived EVs partially rescued the tumor size and weight of the gene-depleted PC3M xenografts. SHC4, PFDN4 and CHORDC1 are expected to be beneficial for the inhibition of cell proliferation and cell-to-cell communication via EVs in PCa.

In summary, our study extensively screened miRNAs that regulate exosome secretion in PCa and found that miR-26a regulates EV secretion by targeting SHC4, PFDN4 and CHORDC1. This finding reveals novel insight into miRNA-mediated tumor suppression by inhibiting EV biogenesis, which may provide novel approaches for PCa treatment.

## Materials and Methods

### Reagents

The following antibodies were used as primary antibodies in immunoblotting: mouse monoclonal anti-human CD9 antibody (clone 12A12, dilution 1:1000) and CD63 antibody (clone 8A12, dilution 1:1000) from Cosmo Bio (Tokyo, Japan). Rabbit polyclonal anti-human PFDN4 (16045-1-AP, dilution 1:200) was from the Proteintech Group. Mouse monoclonal anti-human SHC4 (clone 2F5, dilution 1:300) was from Sigma-Aldrich. Rabbit polyclonal anti-human CHORDC1 (NBP1-78304, dilution 1:2500) was purchased from Novus Biologicals (Littleton, CO). Secondary antibodies (horseradish peroxidase-linked anti-mouse IgG, NA931, or horseradish peroxidase-linked anti-rabbit IgG, NA934, dilution 1:5000) were purchased from GE Healthcare.

The following antibodies were used for ExoScreen analysis: mouse monoclonal anti-human CD9 antibody (clone 12A12) and CD63 antibody (clone 8A12). These antibodies were used to modify either acceptor bead or biotin following the manufacturer’s protocol.

The following miRNA mimic or siRNA was used for the transient transfection assay: miR-26a mimic (MC10249), miR-194-5p mimic (MC10004), miRNA Mimic Negative Control #1 (4464058), TSAP6 siRNA (4392422), Rab27a siRNA (S11693), Rab27b siRNA (S11697), and nSMase2 siRNA (S30925) were purchased from Ambion (Austin, TX, USA). ALL STAR negative control siRNA (SI03650318) and TSG101 siRNA (SI02655184) were purchased from Qiagen (Hilden, Germany). PFDN4 siRNA (siGENOME SMART pool siRNA M-013012), SHC4 siRNA (siGENOME SMART pool siRNA M-031768), CHORDC1 siRNA (siGENOME SMART pool M-019998) and PRKCD siRNA (siGENOME SMART pool M-003524) were purchased from Dharmacon (Lafayette, CO, USA).

### Cell culture

The human prostate cancer epithelial metastatic cell line PC3 (ATCC CRL-1435) was purchased from ATCC. PC3 and PC-3M-luc-C6 (PC3M) (Xenogen, Alameda, CA) were cultured in RPMI 1640 medium (Gibco) supplemented with 10% heat-inactivated fetal bovine serum (FBS) and an antibiotic-antimycotic (Gibco) solution at 37°C. For routine maintenance, each cell line was grown as a monolayer at 37°C with 5% carbon dioxide and 95% relative humidity (RH).

### Preparation of conditioned media and extracellular vesicles

The cells were washed with phosphate-buffered saline (PBS), and the culture medium was replaced with advanced RPMI 1640 medium (Gibco) for PC3M and PC3, containing an antibiotic-antimycotic and 2 mM L-glutamine (Gibco) ^33^. EVs from the conditioned medium were isolated by a differential ultracentrifugation protocol, as we previously reported ^34^. Briefly, the conditioned medium was centrifuged at 2,000xg for 10 min to remove contaminating cells. The resulting supernatants were then transferred to fresh tubes and filtered through a 0.22 µm filter (Millipore). The filtered conditioned medium was centrifuged for 70 min at 110,000x g to pellet the enriched EVs (Beckman Coulter, Brea, CA, USA). The pellets were washed with 11 ml of PBS and ultracentrifuged at 110,000xg for another 70 min. The EV pellets were stored in a refrigerator at 4°C until use.

### Immunoblotting

The EV fraction or the transfected PCa cells was measured for protein content using the Micro BCA Protein Assay kit (Thermo Scientific, Wilmington, DE). Equal amounts of protein were loaded onto 4-20% Mini-PROTEAN TGX gels (Bio-Rad, Munich, Germany). Following electrophoresis (100 V, 30 mA), the proteins were transferred to a polyvinylidene difluoride membrane. The membranes were blocked with Blocking One solution (Nacalai Tesque, Kyoto, Japan) and then incubated with primary antibodies. After washing, the membrane was incubated with horseradish peroxidase-conjugated sheep anti-mouse IgG or donkey anti-rabbit IgG. Bound antibodies were visualized by chemiluminescence using ImmunoStar LD (Wako).

### miRNA-based high-throughput screening using ExoScreen and Cell Growth Assay

High-throughput miRNA screening (total 1728 miRNAs) was performed using the AccuTargetTM Human miRNA mimic library constructed based on miRBase ver.21 (CosmoBio, Tokyo, Japan). A 100 µl PC3M cell suspension of 5000 cells/well (in RPMI containing 10% serum without antibiotics) was seeded into 96 wells and incubated for 24 hours. Then, transfections of 10 nM microRNA mimics or siRNAs were accomplished with the DharmaFECT Transfection Reagent 1 (Dharmacon) according to the manufacturer’s protocol. After 24 hours, the medium was changed to Advanced RPMI 1640 medium containing 2 mM L-glutamine without an antibiotic-antimycotic. Forty-eight hours after the medium change, ExoScreen and cell growth assays, which are described below, were performed.

AlphaLISA reagents (Perkin Elmer, Inc., Waltham, MA 02451, USA) consisted of AlphaScreen Streptavidin-coated donor beads (6760002), AlphaLISA Unconjugated-acceptor beads (6062011) and AlphaLISA Universal buffer (AL001F). AlphaLISA assays were performed in 96-well half area white plates (6005560) and read in an EnSpire Alpha 2300 Mutilabel Plate reader (Perkin Elmer, Inc.). A 96-well white plate was filled with 5 µl of EVs or 10 µl of CM, 10 µl of 5 nM biotinylated antibodies, and 10 µl of 50 µg/ml AlphaLISA acceptor bead-conjugated antibodies in the universal buffer. The plate was then incubated for 1 hour at 37°C. After incubation, 25 µl of 80 µg/ml AlphaScreen streptavidin-coated donor beads was added. The reaction mixture was incubated in the dark for another 30 min at 37°C. Then, the plate was read on the EnSpire Alpha 2300 Multilabel Plate reader using an excitation wavelength of 680 nm and emission detection set at 615 nm. Background signals obtained from filtrated Advanced RPMI or PBS were subtracted from the measured signals. After collecting 10 µl of CM from the 96-well plate, the cellular viability was determined by a colorimetric assay using a Cell Counting Kit-8 (Dojindo Molecular Technologies, Inc.). Plates were read with a spectrophotometer at a wavelength of 450 nm.

### miRNA and siRNA Transient Transfection Assay

A 2 mL PC3M or PC3 cell suspension of 1.0×10^5^ cells/well was seeded into 6-well plates and incubated for 24 hours. Then, transfections of 10 nM microRNA mimics or siRNAs were accomplished with the DharmaFECT Transfection Reagent 1 according to the manufacturer’s protocol. The miR-26a mimic or negative control miRNA (miRNA mimic Negative Control #1, Ambion) was used at a final concentration of 10 nM to investigate the effect of miR-26a on EV secretion. PFDN4 siRNA, CHORDC1 siRNA, SHC4 siRNA, PRKCD siRNA or ALL STAR negative control siRNA was used at a final concentration of 10 nM to investigate the effect of these genes on EV secretion. After 24 hours, the conditioned medium was changed to Advanced RPMI medium containing an antibiotic-antimycotic and 2 mM L-glutamine. Forty-eight hours after the medium change, total RNA was extracted using a miRNeasy Mini Kit (Qiagen) according to the manufacturer’s instructions, and PFDN4, CHORDC1 and SHC4 expression were then determined by qPCR. The CM was collected and purified for EVs by ultracentrifugation.

### Analysis of extracellular vesicle particles by nanoparticle tracking analysis

Isolated EV suspensions diluted in PBS were analyzed by NanoSight particle tracking analysis (LM10, with software version 2.03). For particle tracking, at least five 60 sec videos were taken of each sample with a camera gain of 7. The analysis settings were optimized and kept constant between samples. EV concentrations were calculated as particle/cell of culture to obtain net vesicle secretion rates.

### Microarray and bioinformatics

To perform a mRNA microarray, 1.0×10^5^ PC3M and PC3 cells were seeded into 6-well plates, and 24 hours later, the transfection of miR-26a mimic or controls was accomplished with the DharmaFECT Transfection Reagent 1 (Dharmacon) according to the manufacturer’s protocol. After 24 hours, the CM was changed to Advanced RPMI medium containing 2 mM L-glutamine without an antibiotic-antimycotic. Total mRNAs were extracted from the PC3M and PC3 cells 48 hours after the medium change.

Total RNA was amplified and labelled with Cy3 using a Low Input Quick Amp Labeling Kit, one color (Agilent Technologies), following the manufacturer’s instructions. Briefly, 100 ng of total RNA was reverse-transcribed to double-stranded cDNA using a poly dT-T7 promoter primer. Primer, template RNA and quality-control transcripts of known concentration and quality were first denatured at 65 °C for 10 min and incubated for 2 h at 40 °C with 5 × first-strand buffer, 0.1 M dithiothreitol, 10 mM deoxynucleotide triphosphate mix and AffinityScript RNase Block Mix. The AffinityScript enzyme was inactivated at 70 °C for 15 min. cDNA products were then used as templates for in vitro transcription to generate fluorescent complementary RNA (cRNA). cDNA products were mixed with a transcription master mix in the presence of T7 RNA polymerase and Cy3-labeled CTP (cytidine 5′-triphosphate) and incubated at 40 °C for 2 h. Labeled cRNA was purified using RNeasy Mini Spin Columns (Qiagen) and eluted in 30 μl of nuclease-free water. After amplification and labeling, cRNA quantity and cyanine incorporation were determined using a NanoDrop ND-1000 spectrophotometer and an Agilent Bioanalyzer. For each hybridization, 0.60 μg of Cy3-labelled cRNA was fragmented and hybridized at 65 °C for 17 h to an Agilent SurePrint G3 Human GE v3 8×60K Microarray (design ID: 072363). After washing, the microarray chips were scanned using an Agilent DNA microarray scanner. The intensity values of each scanned feature were quantified using Agilent Feature Extraction software version 11.5.1.1, which performs background subtractions. We only used features that were flagged as no errors (detected flags) and excluded features that were not positive, not significant, not uniform, not above background, saturated and population outliers (compromised and not detected flags). Normalization was performed with Agilent GeneSpring version 14.9.1 (per chip: normalization to 75th percentile shift). There are a total of 58,201 probes on the Agilent SurePrint G3 Human GE v3 8×60K Microarray (design ID: 072363) without control probes. The altered transcripts were quantified using the comparative method.

Unsupervised clustering and heat map generation using Pearson’s correlation in Ward’s method for linkage analysis and principal component analysis (PCA) were performed using Partek Genomics Suite 6.6. GSEA (www.broadinstitute.org/gsea) to compare the gene expression of PC3M and PC3 after transfection of miR-26a with those of the control (miRNA mimic Negative Control #1).

### RNA extraction and qPCR analysis

Total RNA was extracted from cultured cells using QIAzol and the miRNeasy Mini Kit (Qiagen, Hilden, Germany) according to the manufacturer’s protocols. The purity and concentration of all RNA samples were quantified using a NanoDrop ND-1000 spectrophotometer (Thermo Scientific). The reverse transcription reaction was performed using a High-Capacity cDNA Reverse Transcription Kit (Applied Biosystems, Foster City, CA) and a random hexamer primer. Real-time PCR analyses were performed using StepOne Plus and TaqMan Universal PCR Master Mix (Thermo Fisher Scientific). Expression of mRNA was normalized to β-actin. TaqMan probes for PFDN4, SHC4, CHORDC1 and β-actin were purchased from Applied Biosystems.

### Construction of shRNA vector and establishment of stable cell lines

Knockdown vectors expressing shRNAs were constructed by subcloning an annealed oligonucleotide into the pBAsi-hU6-Pur vector (TaKaRa Bio). Oligonucleotide sequences encoding the shRNA hairpin are shown below.

CHORDC1_sense: 5’-GATCCGAAGCAAATAGCACATTGTTAAATCTGTGAAGCCACAGATGGGATTTAACAATGTGCTATTTGCTTCTTTTTTA-3’

CHORDC1_antisense: 5’-AGCTTAAAAAAGAAGCAAATAGCACATTGTTAAATCCCATCTGTGGCTTCACAGATTTAACAATGTGCTATTTGCTTCG-3’

PFDN4_sense: 5’- GATCCGAAGAAATTGACGCCTTAGAATCCCTGTGAAGCCACAGATGGGGGATTCTAAGGCGTCAATTTCTTCTTTTTTA-3’

PFDN4_antisense: 5’-AGCTTAAAAAAGAAGAAATTGACGCCTTAGAATCCCCCATCTGTGGCTTCACAGGGATTCTAAGGCGTCAATTTCTTCG-3’

SHC4_sense: 5’- GATCCGCCCCAGAAACCAGTTTAAGTAGGCTGTGAAGCCACAGATGGGCCTACTTAAACTGGTTTCTGGGGCTTTTTTA-3’

SHC4_antisense:5’- AGCTTAAAAAAGCCCCAGAAACCAGTTTAAGTAGGCCCATCTGTGGCTTCACAGCCTACTTAAACTGGTTTCTGGGGCG-3’).

HindIII (TOYOBO # HND-311) and BamHI (TOYOBO # BAH-111) were used to digest the pBAsi-hU6-Pur vector and the double-stranded shRNA oligonucleotide cassette insert followed by ligation with T4 DNA ligase. The template oligos were purchased from Thermo Fisher Scientific. Inserted sequences were confirmed by Sanger sequencing.

Stable PC3M SHC4-, PFDN4-, or CHORDC1-modified cell lines that expressed shRNA against each human gene were generated by selection with puromycin (2 µg/mL). PC3M cells at 90% confluency were transfected with 0.5 µg of the vector in 24-well dishes using the Lipofectamine LTX Reagent in accordance with the manufacturer’s instructions (Life Technologies). Cells were replaced in a 15-cm dish 12 hours after transfection, followed by selection with puromycin for 2 weeks. Five surviving single colonies were picked from each transfected and cultured for an additional 2 weeks. The cells secreting fewer EVs among the transfectants were considered to be stable cell lines.

### Plasmid constructs and luciferase reporter assay

The following annealed oligos (Thermo Fisher) were used for constructing 3’UTR-reporter vectors.

PFDN4_3UTR_S: 5′- TCGAGACATTTTATAATACTTTTTTTATTTGTTTAATAAACTTGAATATTGTTTAAAATGATAATTTTCCTTCTTCAAATGACATGGAGC-3′

PFDN4_3UTR_AS: 5′- GGCCGCTCCATGTCATTTGAAGAAGGAAAATTATCATTTTAAACAATATTCAAGTTTATTAAACAAATAAAAAAAGTATTATAAAATGTC-3′

SHC4_3UTR_S: 5′- TCGAGAAAAGCACAACTAAAATTTCACATGCTAACGACAACTTGAATGAACTGCTGGGGCAGTGGTATGTGCCTTTCAACTTGATAATTGC-3′

SHC4_3UTR_AS: 5′- GGCCGCAATTATCAAGTTGAAAGGCACATACCACTGCCCCAGCAGTTCATTCAAGTTGTCGTTAGCATGTGAAATTTTAGTTGTGCTTTTC-3′

CHORDC1_1_3UTR_S: 5′- TCGAGTTCTCCCTACTGGTAGGAACCATAGTTGTGTCCTATACTTGAAGAGGCTGGAAAGTAGCCCATAACCATAATTGCAGTATTTCTTGC-3′

CHORDC1_1_3UTR_AS: 5′- GGCCGCAAGAAATACTGCAATTATGGTTATGGGCTACTTTCCAGCCTCTTCAAGTATAGGACACAACTATGGTTCCTACCAGTAGGGAGAAC-3′

The annealed oligo for the 3′UTR of PFDN4, SHC4, or CHORDC1 was subcloned into psiCHECK2 that had been digested with XhoI and NotI. To mutate miR-26a recognition sites in PFDN4, SHC4, or CHORDC1, annealed oligos were used.

PFDN4_3UTR_mut_S: 5′- TCGAGACATTTTATAATACTTTTTTTATTTGTTTAATATTGAACTTTATTGTTTAAAATGATAATTTTCCTTCTTCAAATGACATGGAGC-3′

PFDN4_3UTR_mut_AS: 5′-GGCCGCTCCATGTCATTTGAAGAAGGAAAATTATCATTTTAAACAATAAAGTTCAATATTAAACAAATAAAAAAAGTATTATAAAATGTC-3′

SHC4_3UTR_mut_S: 5′- TCGAGAAAAGCACAACTAAAATTTCACATGCTAACGACATGAACTTTGAACTGCTGGGGCAGTGGTATGTGCCTTTCAACTTGATAATTGC-3′

SHC4_3UTR_mut_AS: 5′- GGCCGCAATTATCAAGTTGAAAGGCACATACCACTGCCCCAGCAGTTCAAAGTTCATGTCGTTAGCATGTGAAATTTTAGTTGTGCTTTTC-3′

CHORDC1_1_3UTR_mut_S: 5′- TCGAGTTCTCCCTACTGGTAGGAACCATAGTTGTGTCCTAATGAACTTGAGGCTGGAAAGTAGCCCATAACCATAATTGCAGTATTTCTTGC-3′

CHORDC1_1_3UTR_mut_AS: 5′- GGCCGCAAGAAATACTGCAATTATGGTTATGGGCTACTTTCCAGCCTCAAGTTCATTAGGACACAACTATGGTTCCTACCAGTAGGGAGAAC-3′

Luciferase reporter assays were performed by cotransfecting 293 cells with 8 µl of 5 µM miR-26a mimic or negative control mimic with 500 ng of psiCHECK2 reporter plasmids using Lipofectamine 3000 (Thermo Fisher) and performing the Dual-Luciferase Reporter Assay System (Promega) after 48 h.

### Mouse studies

Animal experiments in this study were performed in compliance with the guidelines of the Institute for Laboratory Animal Research, National Cancer Center Research Institute. Five-week-old male BALB/C nude mice (Charles River Laboratories, Kanagawa, Japan) were used for animal experiments. We subcutaneously injected PC3M-control, SHC4-, PFDN4-, or CHORDC1-modified cells (1×10^6^ cells were injected in 50-µl volume Matrigel diluted with PBS) into anesthetized mice. We carefully monitored mice and measured the size of their tumors using a Vernier caliper. For the rescue experiment, PC3M-SHC4, PC3M-PFDN4, or PC3M-CHORDC1-KD cells (1×10^6^ cells suspended in 50 µl volume Matrigel diluted with PBS) were subcutaneously injected. After 7 days of implantation, 3 µg of EVs from PC3M control cells was injected intratumorally (50 µl in PBS) every other day for up to 28 days. Mice were monitored carefully, and the size of their tumors was measured using a Vernier caliper. Tumors were harvested 31 days after inoculation of cancer cells, and tumor weight was measured.

### Statistical analysis

Unless otherwise described, the data are presented as the mean±SE, and statistical significance was determined by Student’s t-test. In the dot plot, the bars indicate the median and interquartile range, and statistical significance was determined by Student’s *t*-test. P<0.05 was considered statistically significant.

## Acknowledgments

We thank Ayako Inoue for support on this work. This study was supported by the Practical Research for Innovative Cancer Control (18ck0106366h0001) from the Japan Agency for Medical Research and Development, AMED; and the ‘Development of Diagnostic Technology for Detection of miRNA in Body Fluids’ grant from AMED.

## Supplementary Figure legends

**Supplementary Figure 1.**

A. Correlation matrix between the controls: Positive correlations are shown in red, and negative correlations are shown in blue. The values are the mean±SE (n=18). **, p<0.01; and n.s., not significant.

B. The results of the screening from candidate 30 miRNAs. The effect of 30 miRNAs and nonspecific miRNA mimic (control) on the secretion of EVs and cell viability. The secretion of EV was evaluated by ExoScreen, and the cell viability was measured by the MTS assay.

**Supplementary Figure 2.**

A. A principal component analysis (PCA) map for 99 PCa tissues and 28 normal adjacent benign prostate tissues with 373 miRNAs.

B. Heat map showing the differences in 59 miRNAs whose expression levels were repressed >1.25-fold in prostate cancer tissue relative to normal adjacent benign prostate tissue and p-value <0.001.

C. The effect of the miR-26a mimic on EV secretion per PC3 cell. The secretion of EVs per cell was evaluated by the signal intensity of ExoScreen per cell. The values are depicted as the fold change relative to the nonspecific miRNA mimic (control). The values are the mean±SE (n=3). **, p<0.01.

D. The effect of the miR-26a mimic on EV secretion per PC3 cell. The amount of EV secreted per cell was evaluated using a nanoparticle tracking system. The values are the mean±SE (n=3). **, p<0.01.

**Supplementary Figure 3.**

The effect of candidate gene siRNAs and negative control siRNA (control) on the secretion of EVs and cell viability. The secretion of EV was evaluated by ExoScreen, and the cell viability was measured by the MTS assay. A total of 88 candidate genes were separated into two plates, plate 1 and plate 2.

**Supplementary Figure 4.**

A. The effect of the miR-26a mimic on EV secretion per PC3 cell. The secretion of EVs was evaluated by the signal intensity of ExoScreen. The values are depicted as the fold change relative to the nonspecific miRNA mimic (control). The values are the mean±SE (n=3). **, p<0.01; and n.s., not significant.

B. The effect of the miR-26a mimic on EV secretion per PC3 cell. The particle number of EVs was measured using a nanoparticle tracking system. The values are the mean±SE (n=3). *, p<0.05; and n.s., not significant.

**Supplementary Figure 5.**

A. The effect of miR-26a on the expression level of target genes in PC3M cells. The values are depicted as the fold change relative to the nonspecific miRNA mimic (control). The values are the mean±SE (n=3). **, p<0.01.

B. The effect of miR-26a on the expression level of target genes in PC3M cells. The values are depicted as the fold change relative to the nonspecific miRNA mimic (control). The values are the mean±SE (n=3). **, p<0.01.

**Supplementary Figure 6.**

The xenografts from nude mice injected with PBS or EVs. The tumor weights in nude mice at 35 days were determined. The values are the mean±SE (n=6). *, p<0.05.

