## Supplemental figures for "miR-26a regulates extracellular vesicle secretion from prostate cancer cells via targeting SHC4, PFDN4 and CHORDC1"

### Supplementary Figure 1.

*Urabe F et al.*

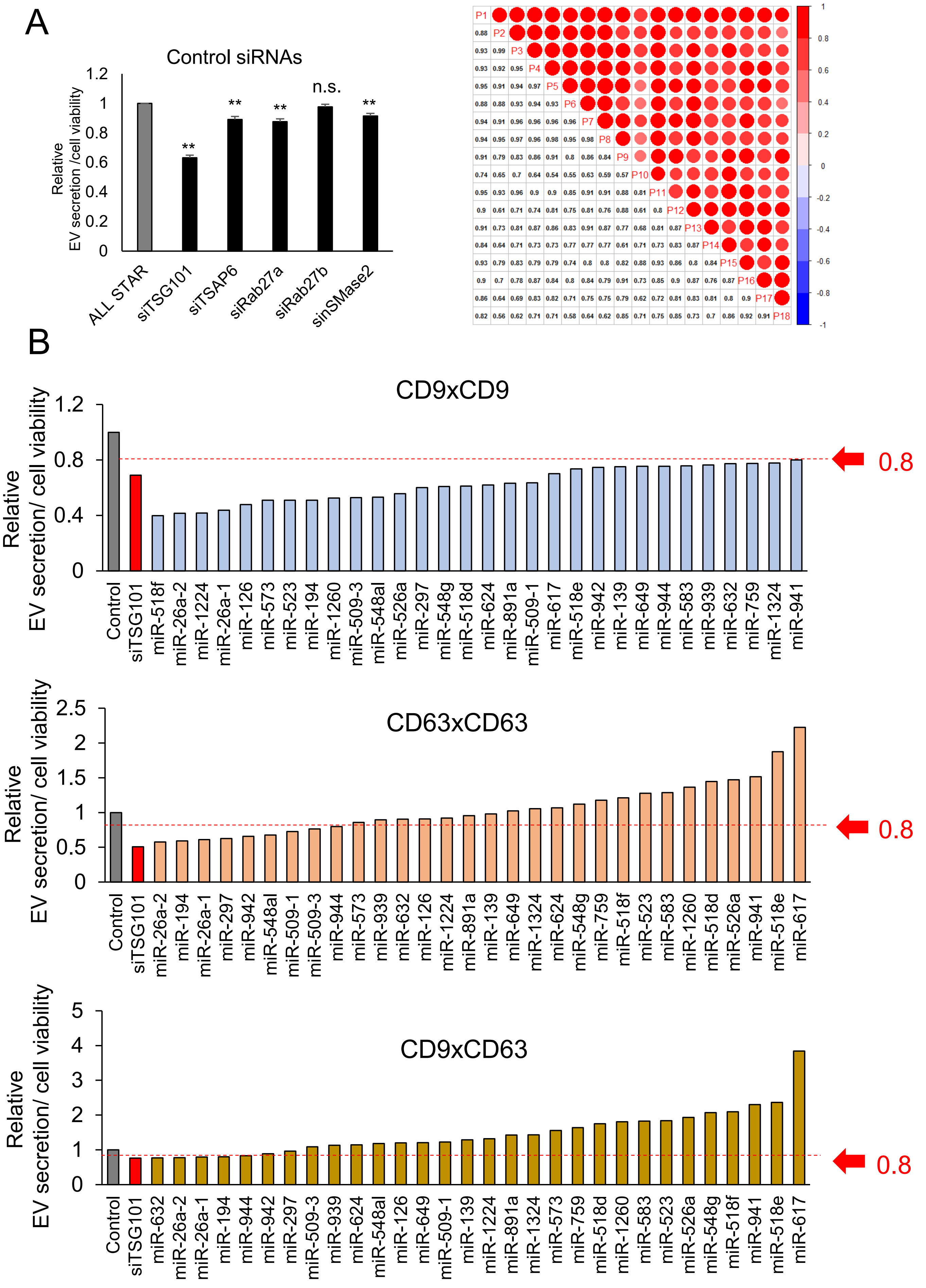

### Supplementary Figure 2.

*Urabe F et al.*

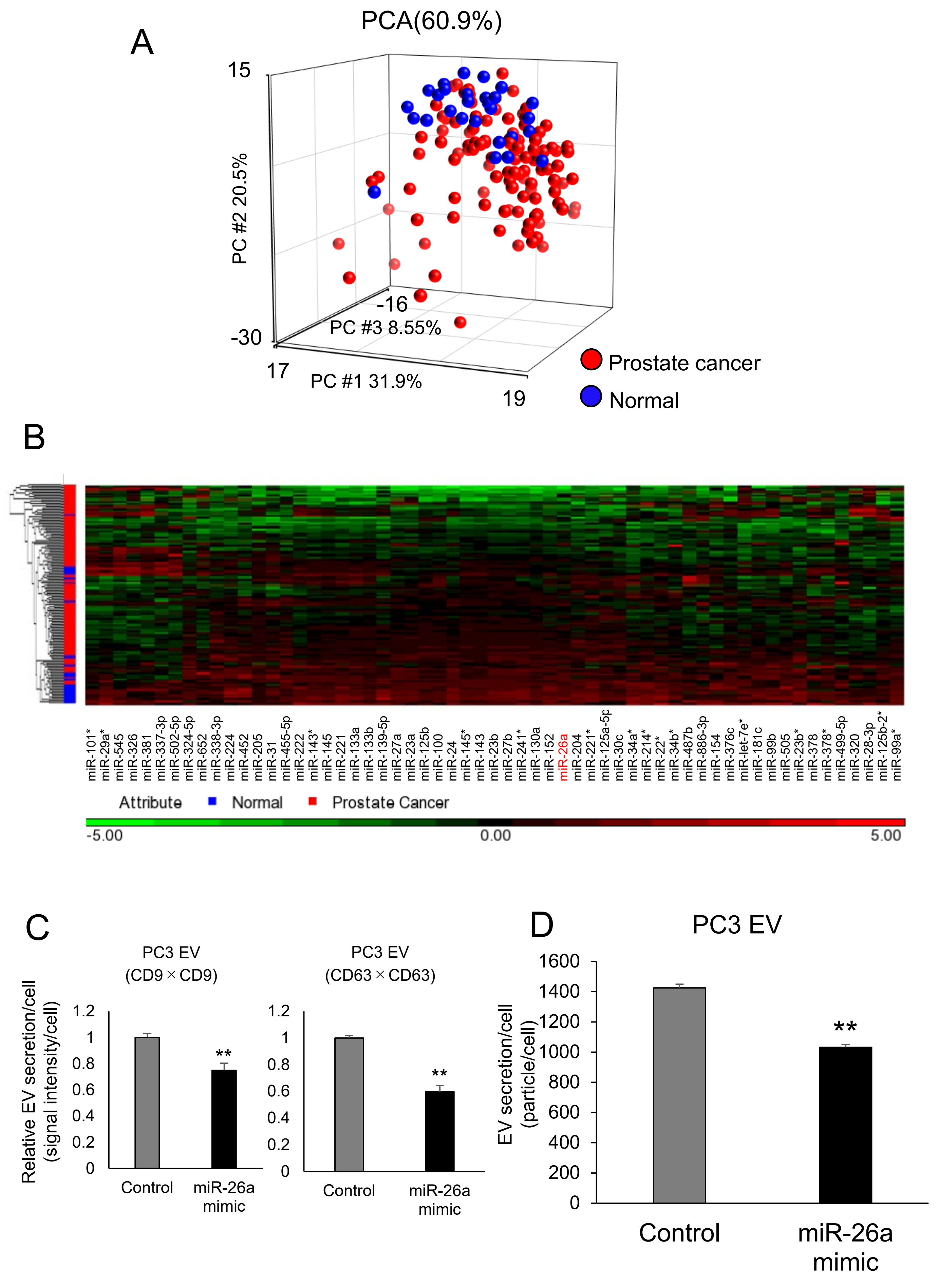

### Supplementary Figure 3.

*Urabe F et al.*

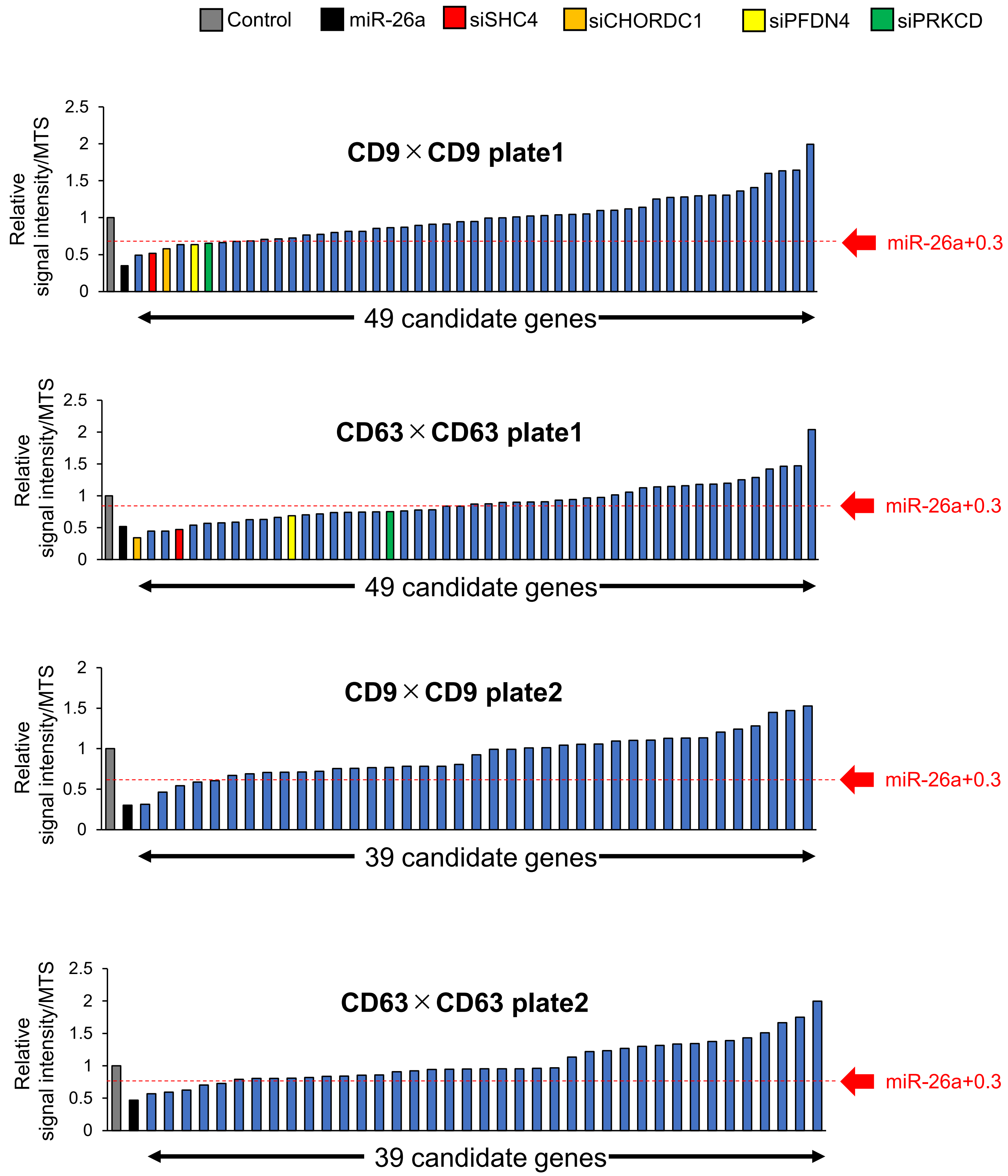

### Supplementary Figure 4.

*Urabe F et al.*

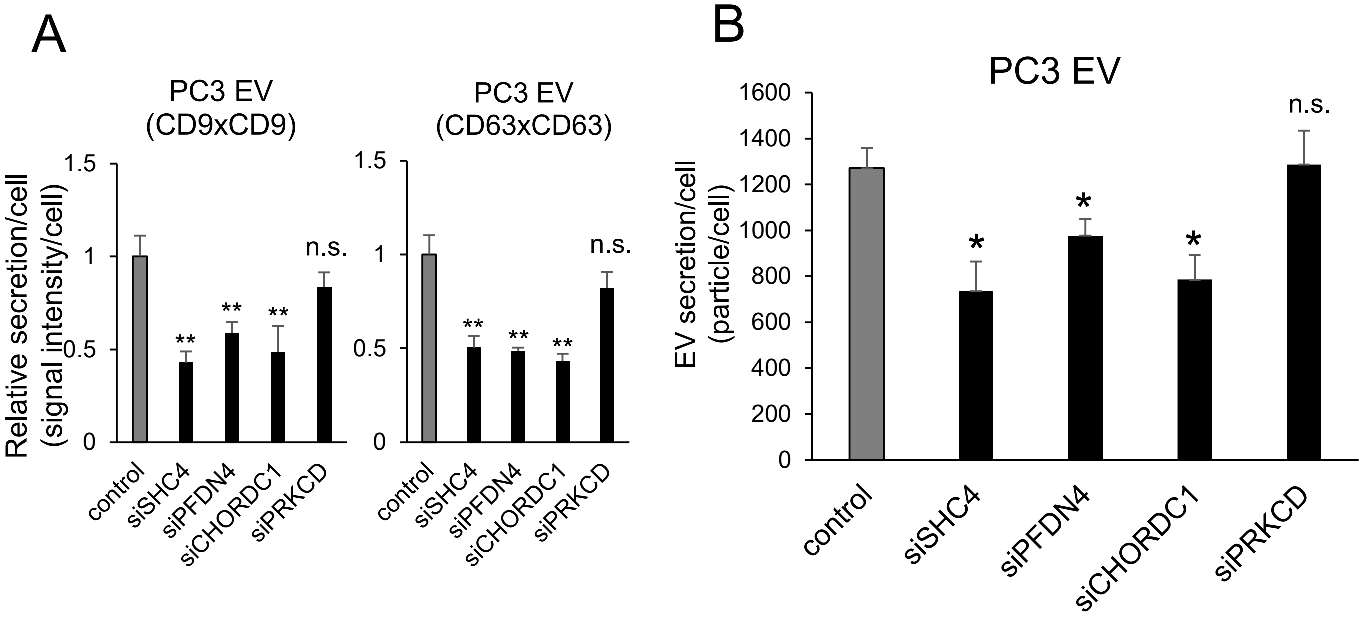

### Supplementary Figure 5.

*Urabe F et al.*

A

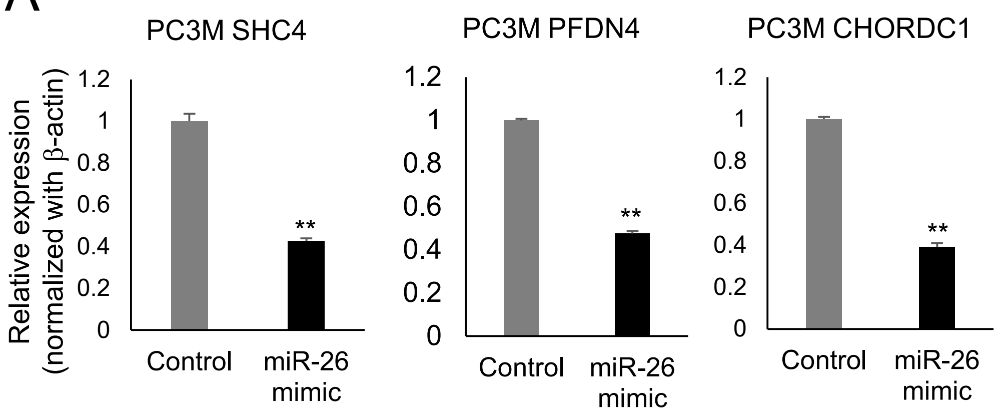


B

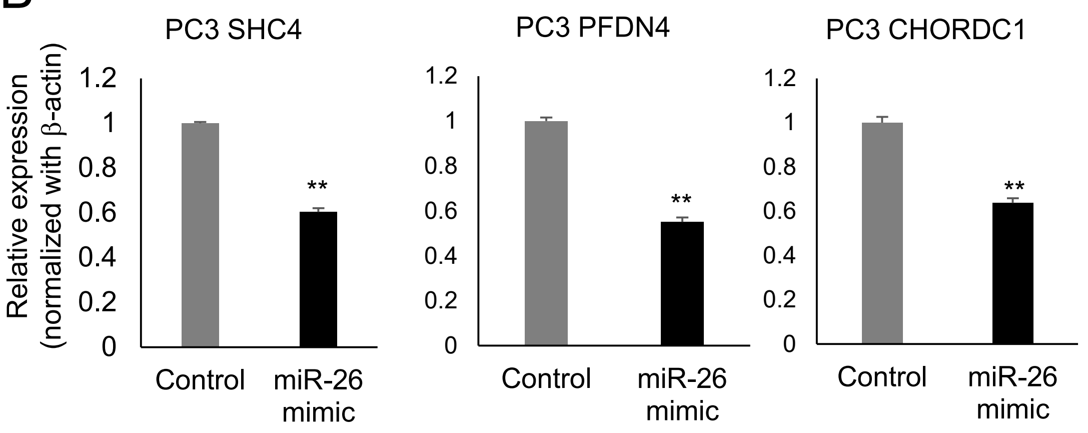

**Supplementary Figure 6.** *Urabe F et al.*

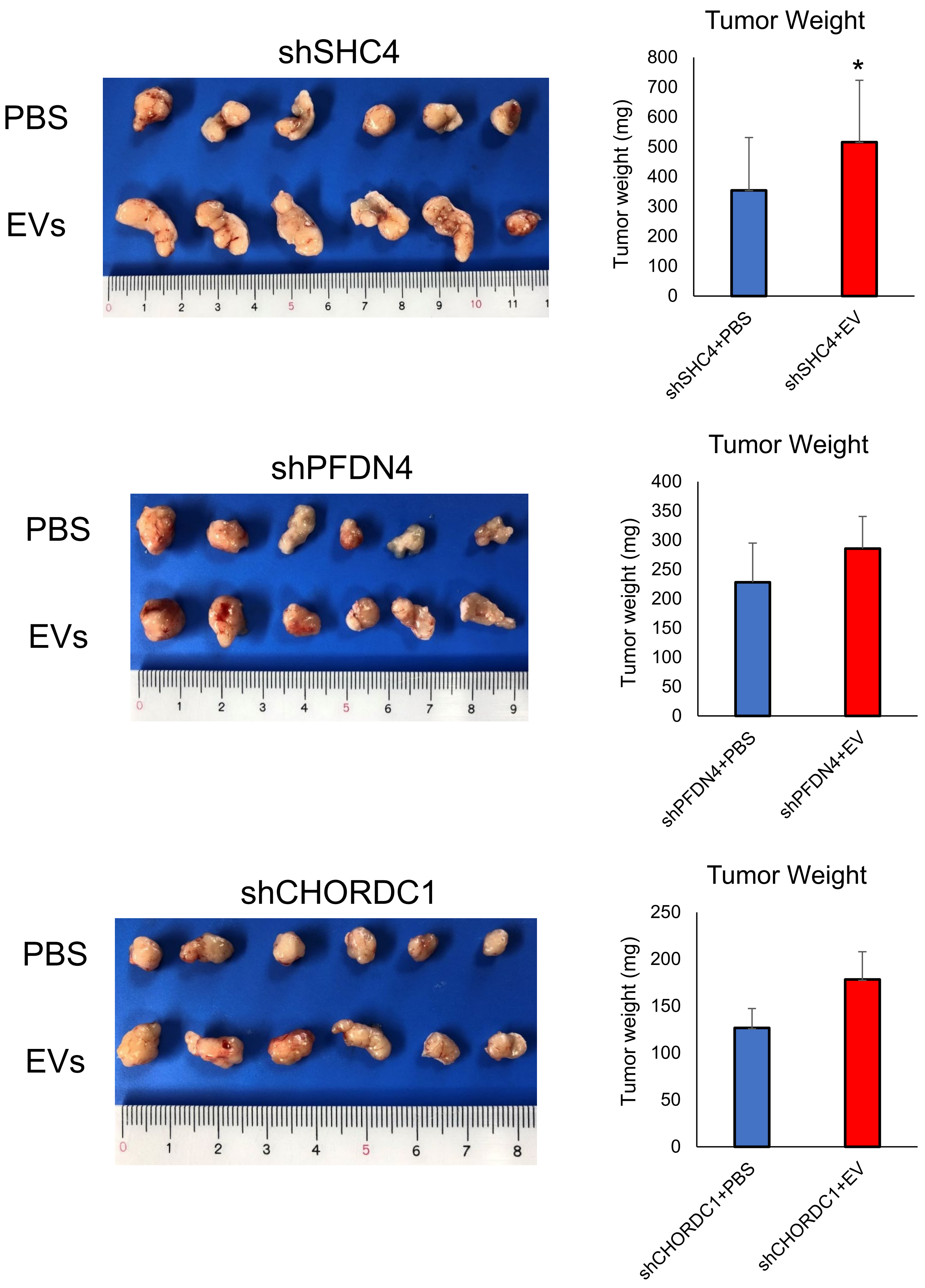
